# *F*_*ST*_ and the Triangle Inequality for Biallelic Markers

**DOI:** 10.1101/567743

**Authors:** Ilana M. Arbisser, Noah A. Rosenberg

## Abstract

The population differentiation statistic *F*_*ST*_, introduced by Sewall Wright, is often treated as a pairwise distance measure between populations. As was known to Wright, however, *F*_*ST*_ is not a true metric because allele frequencies exist for which it does not satisfy the triangle inequality. We prove that a stronger result holds: for biallelic markers whose allele frequencies differ across three populations, *F_ST_ never* satisfies the triangle inequality. We study the deviation from the triangle inequality as a function of the allele frequencies of three populations, identifying frequency vectors at which the deviation is maximal. We also examine the implications of the failure of the triangle inequality for the four-point condition for groups of four populations. Next, we examine the extent to which *F*_*ST*_ fails to satisfy the triangle inequality in genome-wide data from human populations, finding that some loci have frequencies that produce deviations near the maximum. We discuss the consequences of the theoretical results for various types of data analysis, including multidimensional scaling and inference of neighbor-joining trees from pairwise *F*_*ST*_ matrices.

## 1. Introduction

Introduced by Wright (1951), *F*_*ST*_, which provides a measure of population structure for population-genetic data, is one of the most commonly used statistics in population genetics (Holsinger and Weir 2009). Pairwise *F*_*ST*_ computed between two populations is often viewed as a measure of genetic “distance” between the populations (e.g. Jorde 1985, Rosenberg et al. 2005). Indeed, *F*_*ST*_ is frequently treated as a distance in many types of analysis for representing relationships between multiple populations, such as in the distance matrices used for spatially depicting genetic variation and in inference of population trees (e.g. Pérez-Lezaun et al. 1997, Li et al. 2008).

In the formulation of Nei (1973), *F*_*ST*_ can be written

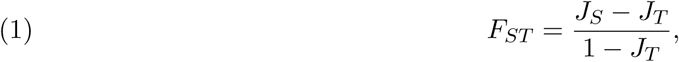

where *J*_*S*_ is the mean homozygosity across a set of subpopulations and *J*_*T*_ is the homozygosity of a population formed by pooling the subpopulations, assuming they have equal representation.

For the case of two subpopulations, using *p*_*ki*_ for the frequency of allele *i* in subpopulation *k*, 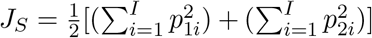 and 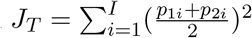, where *I* is the total number of alleles at a locus of interest. Hence, eq. (1) reduces in this case to

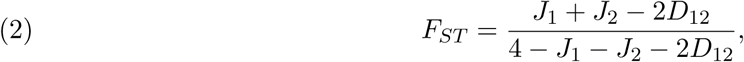

where 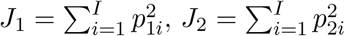, and 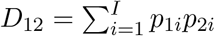.

*F*_*ST*_, eq. (2), has some of the properties required by a mathematical measure of distance: it is symmetric with respect to a change in the population labels, it is nonnegative, and it is equal to 0 if and only if two populations have the same allele frequencies (*p*_1*i*_ = *p*_2*i*_ for all *i*). Yet, *F*_*ST*_ is not a true distance metric because it does not satisfy the triangle inequality: with three populations, the sum of two of the distances can be smaller than the third distance. In fact, Sewall Wright (1978, p. 89) was aware of this fact, offering a counterexample of three populations whose allele frequencies result in values of *F*_*ST*_ that do not satisfy the inequality: a biallelic locus that is monomorphic for one allele in population 1 and monomorphic for the other allele in population 2, and has equal frequencies for the two alleles in population 3.

Here, we generalize beyond Wright’s counterexample to show that not only is it possible for *F*_*ST*_ to violate the triangle inequality, for a biallelic locus with distinct allele frequencies in three populations, *F_ST_ never* satisfies the triangle inequality. We explore the extent to which *F*_*ST*_ fails to satisfy the triangle inequality over the space of possible allele frequencies, finding that the maximal deviation from the condition specified by the triangle inequality occurs precisely in Wright’s counterexample. We also show that the failure to satisfy the triangle inequality has as a consequence a failure of the four-point condition associated with construction of evolutionary trees. To consider the context of our theoretical results in data analysis, we examine the extent to which *F*_*ST*_ fails to satisfy the triangle inequality in data from three human populations. We also examine the impact of the mathematical results in multidimensional scaling analysis and on inference of population trees by neighbor-joining.

## 2. The Triangle Inequality Never Holds for Biallelic Markers with Distinct Allele Frequencies

We consider a biallelic locus in three populations. We choose one allele and label its frequencies in populations 1, 2, and 3, by *p*_1_, *p*_2_, and *p*_3_, respectively. Without loss of generality, we assume 0 ⩽ *p*_1_ ⩽ *p*_2_ ⩽ *p*_3_ ⩽ 1. We can define *F* (*p_i_, p*_*j*_) as the value of *F*_*ST*_ measured between two populations *i* and *j*, in which the frequencies of the chosen allele are *p*_*i*_ and *p*_*j*_, respectively.

Simplifying the expression for *F_ST_* from eq. (2) by noting that the other allele of the biallelic locus has frequency *q_i_* = 1 − *p*_*i*_ and *q*_*j*_ = 1 − *p*_*j*_ in populations *i* and *j*, respectively, *F*_*ST*_ (*p_i_, p*_*j*_), or *F*_*ij*_ for short, can be written

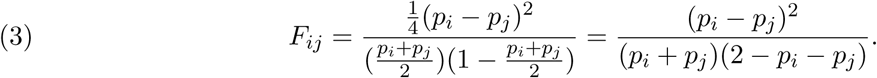

At *p*_*i*_ = *p*_*j*_ = 0 and at *p*_*i*_ = *p*_*j*_ = 1, we define *F*_*ij*_ to be 0. We disregard the cases of *p*_1_ = *p*_2_ = *p*_3_ = 0 and *p*_1_ = *p*_2_ = *p*_3_ = 1, as these cases do not represent polymorphic loci.

The triangle inequality holds for *F*_*ST*_ in three populations if and only if all three of the following inequalities hold:

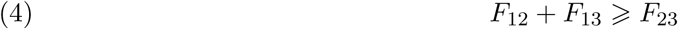

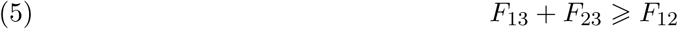

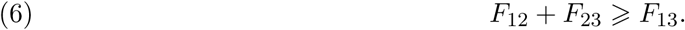

We show that eqs. (4) and (5) always hold, as these statements place the largest of the three *F*_*ST*_ values, *F*_13_, on the larger side of the inequality. We also show that when *p*_1_ = *p*_2_ ⩽ *p*_3_ or *p*_1_ ⩽ *p*_2_ = *p*_3_, eq. (6) also holds, so that the triangle inequality is satisfied. However, we show that when *p*_1_ < *p*_2_ < *p*_3_, the triangle inequality fails: while eqs. (4) and (5) do hold, eq. (6) does not.

### 2.1. At least two of the three of the inequalities always hold

To show that eqs. (4) and (5) hold, it suffices to show that

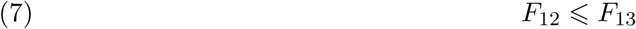

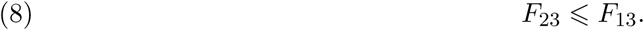

We need only show eq. (7) to also prove eq. (8). In particular, because *p_i_* = 1 − *q_i_* and *p*_1_ ⩽ *p*_2_ ⩽ *p*_3_, *q*_3_ ⩽ *q*_2_ ⩽ *q*_1_ and *F*_*ST*_ (*p_i_, p*_*j*_) = *F*_*ST*_ (*q_i_, q*_*j*_). Hence, by switching the population labels for populations 1 and 3 in eq. (8) and using frequencies *q*_*i*_ in place of *p*_*i*_, eqs. (7) and (8) are equivalent.

To prove eqs. (4) and eq. (5), it remains to prove eq. (7). By eq. (3), we wish to show:

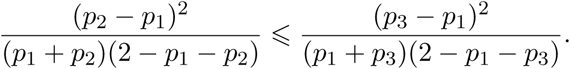

For convenience, we define

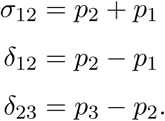

By these definitions,

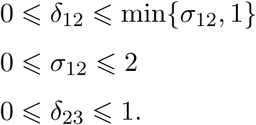

Therefore, we seek to show:

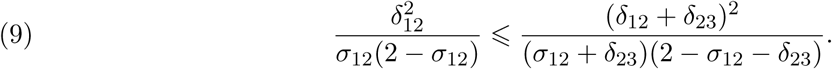

For the cases in which *σ*_12_ = 0 or *σ*_12_ = 2, we have *p*_1_ = *p*_2_ = 0 and *p*_1_ = *p*_2_ = 1, respectively, which we defined in eq. (3) to have *F*_12_ = 0. If *F*_12_ = 0, then *F*_12_ ⩽ *F*_13_ trivially because 0 ⩽ *F*_13_. Hence, noting that *δ*_12_ = 0 if *p*_1_ = *p*_2_, inequality (9) always holds for *p*_1_ = *p*_2_ = *p*_3_.

Similarly, if *σ*_12_ + *δ*_23_ = 0, then *p*_1_ = *p*_3_ = 0, and if *σ*_12_ + *δ*_23_ = 2, then *p*_1_ = *p*_3_ = 1. It follows that *p*_1_ = *p*_3_ = *p*_2_, so the locus is not polymorphic. Consequently, the cases of *σ*_12_ + *δ*_23_ = 0 and *σ*_12_ + *δ*_23_ = 0 are excluded by our assumption that the locus is polymorphic.

Having dealt with the cases in which denominators in eq. (9) are zero, we rearrange eq. (9) and find that eq. (9) holds if

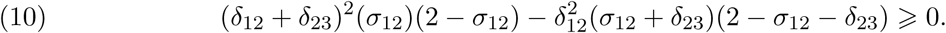

Inequality (10) is equivalent to:

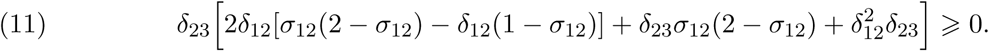

Because *δ*_23_ ⩾ 0, *δ*_23_ does not change the sign of the expression in eq. (11). Hence, eq. (11) holds if *δ*_23_ = 0, or if 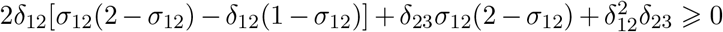. The latter inequality always holds, as 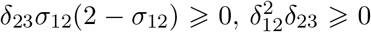, and *σ*_12_(2 − *σ*_12_) − *δ*_12_(1 − *σ*_12_) ⩾ 0, noting that *σ*_12_ ⩾ *δ*_12_, 2 − *σ*_12_ ⩾ 0, and 2 − *σ*_12_ > 1 − *σ*_12_. Therefore, eq. (11) holds, eq. (7) follows, and eqs. (4) and (5) are both true.

### 2.2. If two of the three populations have identical frequencies, then eq. (6) always holds

We show that eq. (6) always holds when *p*_1_ = *p*_2_ ⩽ *p*_3_ or *p*_1_ ⩽ *p*_2_ = *p*_3_. If *p*_1_ = *p*_2_, then *F*_23_ = *F*_13_ and *F*_12_ = 0. Eq. (6) holds trivially as 0+*F*_23_ ⩾ *F*_23_. Similarly, if *p*_2_ = *p*_3_, then *F*_12_ = *F*_13_ and *F*_23_ = 0. Eq. (6) holds trivially, as *F*_12_ + 0 ⩾ *F*_12_. Because eqs. (4)-(6) are all satisfied, the triangle inequality is satisifed for three populations if either *p*_1_ = *p*_2_ ⩽ *p*_3_ or *p*_1_ ⩽ *p*_2_ = *p*_3_.

Note that the triangle inequality is satisfied in the case that two of three points are the same for any function, *f* on some set *X*, which is symmetric and has the identity of indiscernibles, i.e. *f* (*x, x*) = 0 for all *x ∈ X*. With the added condition of nonnegativity, such a function is sometimes called a distance *function* or distance *measure* (to distinguish it from a true distance *metric* that also satisfies the triangle inequality).

### 2.3. If the three populations have distinct frequencies, then eq. (6) never holds

To show that eq. (6) does not hold when 0 ⩽ *p*_1_ < *p*_2_ < *p*_3_ ⩽ 1, we consider a function

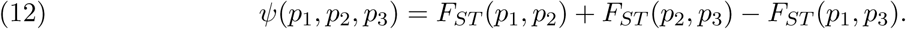

Eq. (6) holds and hence the triangle inequality holds if and only if *ψ*(*p*_1_, *p*_2_, *p*_3_) ⩾ 0. We proceed in several steps to show that *ψ*(*p*_1_, *p*_2_, *p*_3_) < 0 when 0 ⩽ *p*_1_ < *p*_2_ < *p*_3_ ⩽ 1.

i. We write *ψ*(*p*_1_, *p*_2_, *p*_3_) as a fraction with positive denominator and relate *ψ*(*p*_1_, *p*_2_, *p*_3_) to its numerator *ω*(*p*_1_, *p*_2_, *p*_3_). Because the denominator of *ψ*(*p*_1_, *p*_2_, *p*_3_) is always positive, *ψ*(*p*_1_, *p*_2_, *p*_3_) < 0 if and only if *ω*(*p*_1_, *p*_2_, *p*_3_) < 0.
ii. Next, we show that as a function of any one of its variables, *ω*(*p*_1_, *p*_2_, *p*_3_) is a quartic function. Considering *ω*(*p*_1_, *p*_2_, *p*_3_) as a function of *p*_2_, two of its roots lie at *p*_2_ = *p*_1_ and *p*_2_ = *p*_3_. Therefore, we can define a quadratic function in *p*_2_, *φ*(*p*_1_, *p*_2_, *p*_3_):

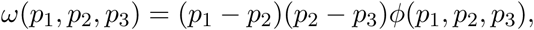 Because we assume *p*_1_ < *p*_2_ and *p*_2_ < *p*_3_, *ω*(*p*_1_, *p*_2_, *p*_3_) < 0 if and only if *φ*(*p*_1_, *p*_2_, *p*_3_) < 0.
iii. We then consider the roots of *φ*(*p*_1_, *p*_2_, *p*_3_) as functions of *p*_2_, *r*_1_(*p*_1_, *p*_3_) and *r*_2_(*p*_1_, *p*_3_). We show that *r*_1_(*p*_1_, *p*_3_) ⩽ *p*_1_, *r*_2_(*p*_1_, *p*_3_) ⩾ *p*_3_, and *φ*(*p*_1_, *p*_2_, *p*_3_) < 0 for *p*_2_ *∈* (*r*_1_(*p*_1_, *p*_3_), *r*_2_(*p*_1_, *p*_3_)).
iv. We conclude that because *φ*(*p*_1_, *p*_2_, *p*_3_) < 0, *ω*(*p*_1_, *p*_2_, *p*_3_) < 0, which implies *ψ*(*p*_1_, *p*_2_, *p*_3_) < 0.

Hence when 0 ⩽ *p*_1_ < *p*_2_ < *p*_3_ ⩽ 1, eq. (6) does not hold, and the triangle inequality does not hold for a biallelic marker with distinct allele frequencies in three populations.

#### 2.3.1. *ψ*(*p*_1_, *p*_2_, *p*_3_) < 0 *if and only if ω*(*p*_1_, *p*_2_, *p*_3_) < 0 *for* 0 ⩽ *p*_1_ < *p*_2_ < *p*_3_ ⩽ 1

We simplify eq. (12) for *ψ*(*p*_1_, *p*_2_, *p*_3_) by using eq. (3)

Define *ω*(*x, y, z*) as:

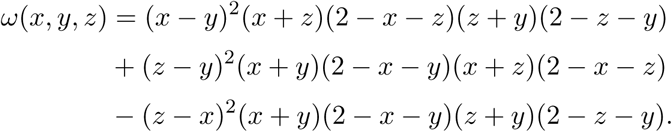

If *x* = *p*_1_, *y* = *p*_2_, and *z* = *p*_3_, then

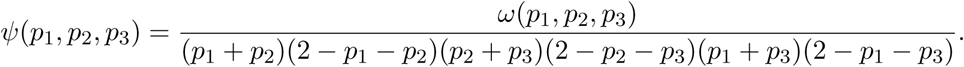

The denominator is always nonnegative when 0 ⩽ *p*_1_ < *p*_2_ < *p*_3_ ⩽ 1, so *ψ*(*p*_1_, *p*_2_, *p*_3_) < 0 if and only if *ω*(*p*_1_, *p*_2_, *p*_3_) < 0.

#### 2.3.2. *ω*(*p*_1_, *p*_2_, *p*_3_) < 0 *if and only if φ*(*p*_1_, *p*_2_, *p*_3_) < 0 *for* 0 ⩽ *p*_1_ < *p*_2_ < *p*_3_ ⩽ 1

Consider *ω*(*x, y, z*) with *x* = *p*_1_ and *z* = *p*_3_ fixed, so that *ω*(*p*_1_, *y, p*_3_) is only a function of *y*. Because *ω* is quartic in *y*, it has at most four distinct roots, each of which can be expressed as a function of *p*_1_ and *p*_3_.

It is trivial to show that *y* = *p*_1_ and *y* = *p*_3_ are both roots of *ω*(*p*_1_, *y, p*_3_). Consequently, *ω*(*p*_1_, *y, p*_3_) has at most two other roots for *y*. Define a quadratic function *φ*(*p*_1_, *y, p*_3_) such that:

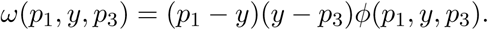

Performing polynomial division, we can write

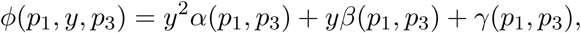

where:

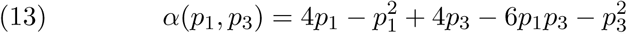

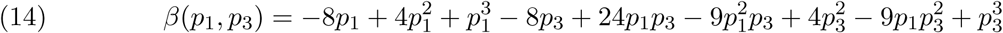

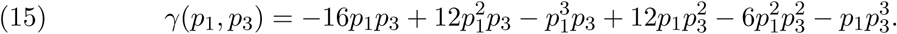

For 0 ⩽ *p*_1_ < *p*_2_ < *p*_3_ ⩽ 1, *p*_1_ − *p*_2_ < 0 and *p*_2_ − *p*_3_ < 0, so *ω*(*p*_1_, *p*_2_, *p*_3_) < 0 if and only if *φ*(*p*_1_, *p*_2_, *p*_3_) < 0.

#### 2.3.3. *φ*(*p*_1_, *p*_2_, *p*_3_) < 0 *when* 0 ⩽ *p*_1_ < *p*_2_ < *p*_3_ ⩽ 1

We know that *φ*(*p*_1_, *y, p*_3_) has at most two roots, *r*_1_(*p*_1_, *p*_3_) and *r*_2_(*p*_1_, *p*_3_). If these two roots are distinct, then *φ*(*p*_1_, *y, p*_3_) > 0 or *φ*(*p*_1_, *y, p*_3_) < 0 for values of *y* between the roots. Without loss of generality, assume *r*_1_(*p*_1_, *p*_3_) ⩽ *r*_2_(*p*_1_, *p*_3_). To show that *φ*(*p*_1_, *p*_2_, *p*_3_) < 0 for 0 ⩽ *p*_1_ < *p*_2_ < *p*_3_ ⩽ 1, we need to show all of the following:

1. If *r*_1_(*p*_1_, *p*_3_) < *y < r*_2_(*p*_1_, *p*_3_), then *φ*(*p*_1_, *y, p*_3_) < 0,
2. *r*_1_(*p*_1_, *p*_3_) ⩽ *p*_1_,
3. *r*_2_(*p*_1_, *p*_3_) ⩾ *p*_3_.

Note that because *p*_1_ < *p*_3_, demonstrating (2) and (3) suffices to show that the two roots *r*_1_(*p*_1_, *p*_3_) and *r*_2_(*p*_1_, *p*_3_) are distinct.

##### 2.3.3.1. *If r*_1_(*p*_1_, *p*_3_) < *y < r*_2_(*p*_1_, *p*_3_), *then φ*(*p*_1_, *y, p*_3_) < 0

*φ*(*p*_1_, *y, p*_3_) is quadratic in *y* with leading coefficient *α*(*p*_1_, *p*_3_). If *α*(*p*_1_, *p*_3_) > 0 and the two roots *r*_1_(*p*_1_, *p*_3_) and *r*_3_(*p*_1_, *p*_3_) are distinct, then *φ*(*p*_1_, *y, p*_3_) < 0 between the roots of *φ*.

To show that *α*(*p*_1_, *p*_3_) > 0, rewrite *α*(*p*_1_, *p*_3_) as follows:

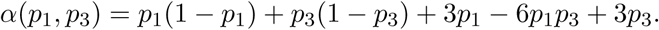

Because 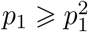 and 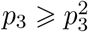, we then have

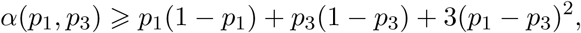

from which we conclude *α*(*p*_1_, *p*_3_) > 0 because *p*_1_ ≠ *p*_3_.

##### 2.3.3.2. *r*_1_(*p*_1_, *p*_3_) ⩽ *p*_1_

Note that *r*_1_(*p*_1_, *p*_3_) ⩽ 0 implies that *r*_1_(*p*_1_, *p*_3_) ⩽ *p*_1_ because 0 ⩽ *p*_1_. We can solve the quadratic equation *φ*(*p*_1_, *y, p*_3_) = 0 for the value of *y* as a function of *p*_1_ and *p*_3_, taking the smaller root to be *r*_1_(*p*_1_, *p*_3_):

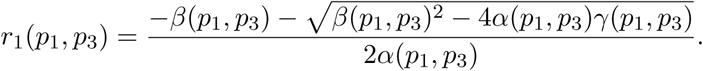

To show *r*_1_(*p*_1_, *p*_3_) ⩽ 0 ⩽ *p*_1_, because we have demonstrated that *α*(*p*_1_, *p*_3_) > 0, we must show

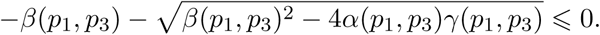

It suffices to show that *γ*(*p*_1_, *p*_3_) ⩽ 0.

Write 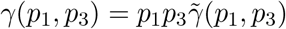, where

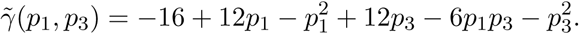

The partial derivatives of 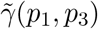 are positive for 0 ⩽ *p*_1_ < *p*_3_ ⩽ 1:

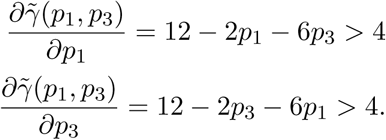

Hence, 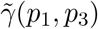 is increasing in *p*_1_ *∈* [0, 1] and *p*_3_ *∈* [*p*_1_, 1] and is maximized at (*p*_1_, *p*_3_) = (1, 1). Because 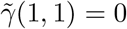, it follows that 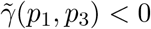 for all (*p*_1_, *p*_3_) with 0 ⩽ *p*_1_ < 1 and *p*_1_ ⩽ *p*_3_ < 1.

We conclude *γ*(*p*_1_, *p*_3_) ⩽ 0 and therefore *r*_1_(*p*_1_, *p*_3_) ⩽ *p*_1_.

#### 2.3.3.3. *r*_2_(*p*_1_, *p*_3_) ⩾ *p*_3_

It suffices to show *r*_2_(*p*_1_, *p*_3_) ⩾ 1 ⩾ *p*_3_.

Taking the positive root of the quadratic equation *φ*(*p*_1_, *p*_2_, *p*_3_) = 0,

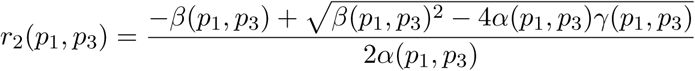

Because *α*(*p*_1_, *p*_3_) > 0 and *γ*(*p*_1_, *p*_3_) ⩽ 0, and leaving off the arguments, it suffices to show *α* + *β* + *γ* ⩽ 0. If *α* + *β* + *γ* ⩽ 0 then 4*α*^2^ + 4*αβ* + 4*αγ* ⩽ 0, (2*α* + *β*)^2^ ⩽ *β*^2^ *−* 4*αγ*, and, thus, 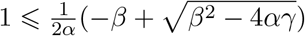.

Define *g*(*p*_1_, *p*_3_) = *α* + *β* + *γ.* We can simplify the condition *g*(*p*_1_, *p*_3_) ⩽ 0, by using eqs. (13)-(15):

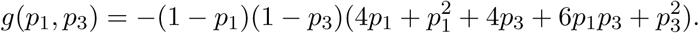

If 0 ⩽ *p*_1_ < *p*_3_ ⩽ 1, then *g*(*p*_1_, *p*_3_) ⩽ 0, as all of the factors in parentheses are nonnegative. Because *g*(*p*_1_, *p*_3_) ⩽ 0, it follows that *r*_2_(*p*_1_, *p*_3_) ⩾ *p*_3_.

We conclude that *φ*(*p*_1_, *p*_2_, *p*_3_) < 0 if 0 ⩽ *p*_1_ < *p*_2_ < *p*_3_ ⩽ 1. We have *r*_1_(*p*_1_, *p*_3_) ⩽ *p*_1_ and *r*_1_(*p*_1_, *p*_3_) ⩾ *p*_3_, the roots of *φ*(*p*_1_, *y, p*_3_) = 0 are distinct, and *φ*(*p*_1_, *y, p*_3_) < 0 between the roots, *r*_1_(*p*_1_, *p*_3_) and *r*_2_(*p*_1_, *p*_3_).

#### 2.3.4. Concluding the proof

Because we have shown that for 0 ⩽ *p*_1_ < *p*_2_ < *p*_3_ ⩽ 1, *φ*(*p*_1_, *p*_2_, *p*_3_) < 0 and *φ*(*p*_1_, *p*_2_, *p*_3_) < 0 implies *ω*(*p*_1_, *p*_2_, *p*_3_) < 0, in turn implying *ψ*(*p*_1_, *p*_2_, *p*_3_) < 0 for 0 ⩽ *p*_1_ < *p*_2_ < *p*_3_ ⩽ 1, eq. (6) is never satisfied, and the triangle inequality is never satsified for biallelic markers with 0 ⩽ *p*_1_ < *p*_2_ < *p*_3_ ⩽ 1.

## 3. The Maximal Deviation from the Triangle Inequality Occurs at Sewall Wright’s Counterexample

### 3.1. Visualization of *ψ*(*p*_1_, *p*_2_, *p*_3_)

As shown in Section 2.3, *ψ*(*p*_1_, *p*_2_, *p*_3_), measuring the extent to which the triangle inequality fails to be satisfied, is always less than or equal to 0 for 0 ⩽ *p*_1_ ⩽ *p*_2_ ⩽ *p*_3_ ⩽ 1. We illustrate the value of *ψ*(*p*_1_, *p*_2_, *p*_3_) over the space of possible allele frequencies (*p*_1_, *p*_2_, *p*_3_) in Figure 1, holding *p*_2_ constant at each of several values and plotting *ψ*(*p*_1_, *p*_2_, *p*_3_) as a function of (*p*_1_, *p*_3_) over the permissible domain [0, *p*_2_] × [*p*_2_, 1].

**Figure 1.**
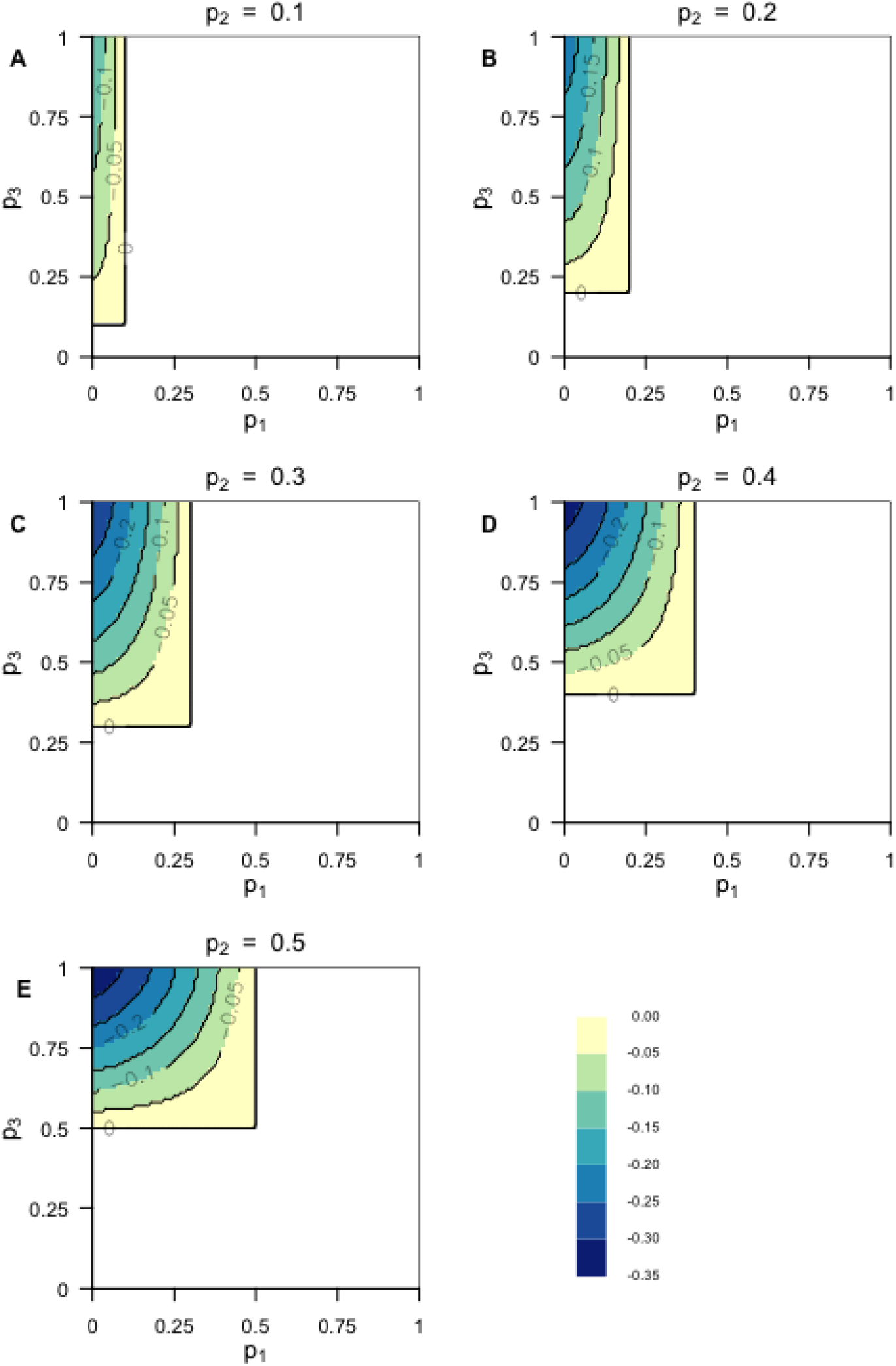
The extent to which the triangle inequality is violated, *ψ*(*p*_1_, *p*_2_, *p*_3_), for constant *p*_2_ (eq. (12)). (A) *p*_2_ = 0.1. (B) *p*_2_ = 0.2. (C) *p*_2_ = 0.3. (D) *p*_2_ = 0.4. (E) *p*_2_ = 0.5. The color at a point represents the value of *ψ* at that point. Because we define *p*_1_ ⩽ *p*_2_ ⩽ *p*_3_, the bottom and right portions of the graph are empty. Because of the symmetry of *ψ* with respect to choice of allele, so that *ψ*(*p*_1_, *p*_2_, *p*_3_) = *ψ*(1 − *p*_3_, 1 − *p*_2_, 1 − *p*_1_) (eq. (19)), we show plots only for *p*_2_ ⩽ 0.5.

In each plot at a fixed *p*_2_, the value of *ψ* appears to decrease monotonically from 0 along lines of constant *p*_1_ and along lines of constant *p*_3_ to a minimum at (*p*_1_, *p*_3_) = (0, 0). Moreover, considering all plots at different values of *p*_2_, the minimum at (*p*_1_, *p*_3_) = (0, 1) appears lowest in the case that 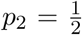. The plots suggest that the point at which *ψ*(*p*_1_, *p*_2_, *p*_3_) is the most negative—where *F*_*ST*_ fails the triangle inequality by the largest amount—is where *p*_1_, *p*_2_, and *p*_3_ are furthest apart. They suggest that the minimum of *ψ*(*p*_1_, *p*_2_, *p*_3_) lies at 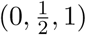, exactly the triplet Sewall Wright (1978, p. 89) offered in his counterexample. We next prove this to be the case.

### 3.2. The minimum of *ψ*(*p*_1_, *p*_2_, *p*_3_) is 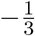 and occurs at 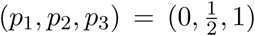

We seek to find the minimum of *ψ*(*p*_1_, *p*_2_, *p*_3_), as described in eq. (12), considering all possible (*p*_1_, *p*_2_, *p*_3_) with 0 ⩽ *p*_1_ ⩽ *p*_2_ ⩽ *p*_3_ ⩽ 1. Note that we can assume 0 ⩽ *p*_1_ < *p*_2_ < *p*_3_ ⩽ 1, because if *p*_1_ = *p*_2_ ⩽ *p*_3_ or *p*_1_ ⩽ *p*_2_ = *p*_3_, then *ψ*(*p*_1_, *p*_2_, *p*_3_) = 0. As we showed previously, *ψ*(*p*_1_, *p*_2_, *p*_3_) = 0 is the maximal value of *ψ*, so in finding the minimum, we can assume *p*_1_, *p*_2_, and *p*_3_ are distinct.

We show that the minimum of *ψ*(*p*_1_, *p*_2_, *p*_3_) occurs at 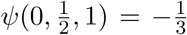. The proof proceeds in three steps:

1. For fixed *p*_2_, *p*_3_, we show *ψ*(0, *p*_2_, *p*_3_) < *ψ*(*p*_1_, *p*_2_, *p*_3_), for all *p*_1_ with 0 < *p*_1_ < 1.
2. For fixed *p*_2_, we show *ψ*(0, *p*_2_, 1) < *ψ*(0, *p*_2_, *p*_3_) for all *p*_3_ with 0 < *p*_3_ < 1.
3. We show 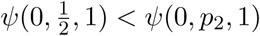 for all *p*_2_ with 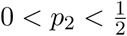 or 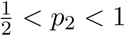.

Showing (1), (2), and (3) suffices to show that the minimum is 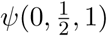, as Step 1 shows that the minimum has *p*_1_ = 0, Step 2 shows that *p*_3_ = 1, and Step 3 shows that 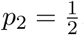.

#### 3.2.1. *ψ*(0, *p*_2_, *p*_3_) < *ψ*(*p*_1_, *p*_2_, *p*_3_)

To show that the minimum of *ψ*(0, *p*_2_, *p*_3_) over 0 ⩽ *p*_1_ ⩽ 1 at fixed (*p*_2_, *p*_3_) occurs at *p*_1_ = 0, we seek to show that there is no minimum of *ψ*(0, *p*_2_, *p*_3_) for 0 < *p*_1_ < 1, so that the minimum must occur on the boundary of the unit interval. If *∂ψ/∂p*_1_ > 0 for 0 < *p*_1_ < 1, then a minimum occurs at the lower bound of *p*_1_: *p*_1_ = 0. To show that *ψ*(0, *p*_2_, *p*_3_) < *ψ*(*p*_1_, *p*_2_, *p*_3_) for all *p*_1_, we show that *∂ψ/∂p*_1_ > 0 everywhere in 0 < *p*_1_ < 1.

Note that

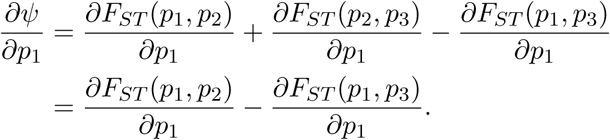

To show *∂F*_*ST*_ (*p*_1_, *p*_2_)*/∂p*_1_*−∂F*_*ST*_ (*p*_1_, *p*_3_)*/∂p*_1_ > 0, it suffices to show *∂F*_*ST*_ (*p*_1_, *p*_2_)*/∂p*_1_ > *∂F*_*ST*_ (*p*_1_, *p*_3_)*/∂p*_1_.

Define a function *f* (*p*_1_, *ρ*) = *∂F*_*ST*_ (*p*_1_, *ρ*)*/∂p*_1_. Note that showing *f* (*p*_1_, *p*_2_) > *f* (*p*_1_, *p*_3_) where *p*_2_ < *p*_3_ is the same as showing that *∂f* (*p*_1_, *ρ*)*/∂ρ* < 0, for 0 < *ρ* < 1 (*ρ* must be strictly in the bounds of its domain because *ρ* = 0 would imply *p*_1_ = *p*_2_ and *ρ* = 1 would imply *p*_2_ = *p*_3_). Showing that *∂*^2^*F*_*ST*_ (*p*_1_, *ρ*)*/∂p*_1_*∂ρ* < 0 implies that *∂ψ*(*p*_1_, *p*_2_, *p*_3_)*/p*_1_ > 0.

Taking the partial derivative of *F*_*ST*_ (*p*_1_, *ρ*) with respect to *p*_1_ gives us

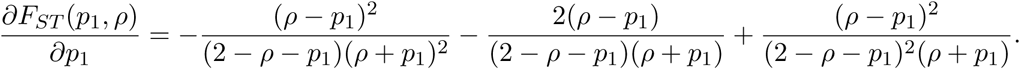

Taking the partial derivative again with respect to *ρ* yields

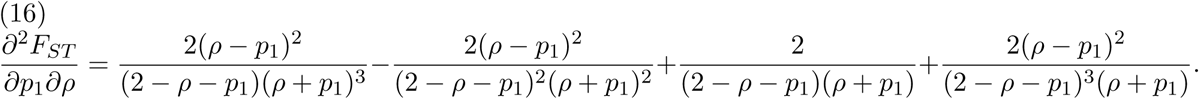

We seek to show that eq. (16) is strictly less than 0. By rearranging terms, it is equivalent to show that

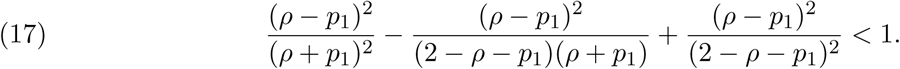

First, consider that 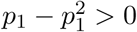 because *p*_1_ < 1 and *ρ − ρ*^2^ > 0 because *ρ* < 1. Thus,

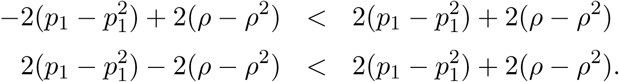

Hence,

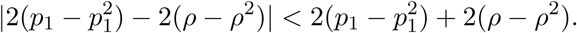

Because *ρ* − *p*_1_, *ρ* + *p*_1_, and 2 − *ρ* − *p*_1_ are positive, we have

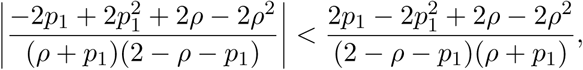

which can be rearranged to show

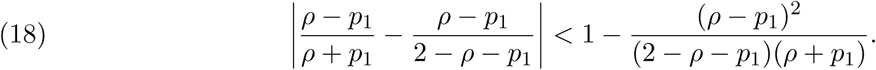

By noting that (*ρ − p*_1_)^2^*/*[(2 − *ρ − p*_1_)(*ρ* + *p*_1_)] is the expression for *F*_*ST*_ (eq. (3)), which is non-negative and less than or equal to 1, we find that both sides of eq. (18) are bounded in [0, 1]:

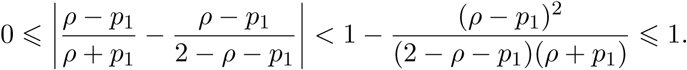

Therefore,

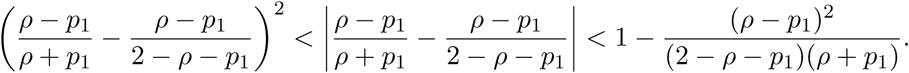

By adding (*ρ − p*_1_)^2^*/*[(2 − *ρ − p*_1_)(*ρ* + *p*_1_)] to both sides of

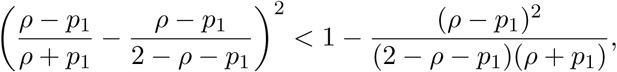

we have eq. (17).

Because eq. (17) holds, we have completed our proof that eq. (16) is less than 0 for 0 < *p*_1_ < 1 and 0 < *ρ* < 1. Therefore, *∂ψ/∂p*_1_ > 0 for 0 < *p*_1_ < 1 at fixed *p*_2_ and *p*_3_, and the minimum of *ψ* occurs at *p*_1_ = 0. We conclude *ψ*(0, *p*_2_, *p*_3_) < *ψ*(*p*_1_, *p*_2_, *p*_3_).

#### 3.2.2. *ψ*(0, *p*_2_, 1) < *ψ*(0, *p*_2_, *p*_3_)

To show *ψ*(0, *p*_2_, 1) < *ψ*(0, *p*_2_, *p*_3_), we first comment that *ψ*(*p*_1_, *p*_2_, *p*_3_) symmetric with respect to the choice of allele, so that

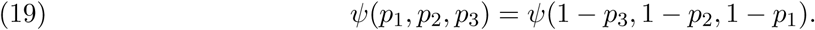

*F*_*ST*_ is symmetric with respect to an exchange of populations: *F*_*ST*_(*p*_*i*_, *p*_*j*_) = *F*_*ST*_(*p*_*j*_, *p*_*i*_). It is also symmetric in the choice of allele used for the computation, so that *F*_*ST*_(*p*_*i*_, *p*_*j*_) = *F*_*ST*_(1 − *p*_*i*_, 1 − *p*_*j*_).

Thus, we have

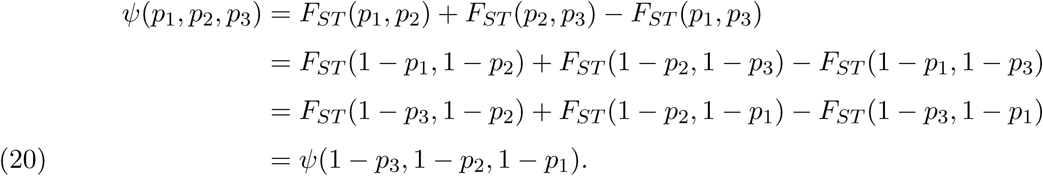

From Section 3.2.1, we have *ψ*(0, *p*_2_, *p*_3_) < *ψ*(*p*_1_, *p*_2_, *p*_3_). By the symmetry in eq. (20), we have *ψ*(1 − *p*_3_, 1 − *p*_2_, 1 *−* 0) < *ψ*(1 − *p*_1_, 1 − *p*_2_, 1 − *p*_3_). Defining *q*_1_ = 1 − *p*_3_, *q*_2_ = 1 − *p*_2_, and *q*_3_ = 1 − *p*_1_, where 0 ⩽ *q*_1_ < *q*_2_ < *q*_3_ ⩽ 1, we can also express this inequality as *ψ*(*q*_1_, *q*_2_, 1) < *ψ*(*q*_1_, *q*_2_, *q*_3_), for all 0 ⩽ *q*_1_ < *q*_2_ < *q*_3_ ⩽ 1. Therefore *p*_1_ = 0 minimizes *ψ* for all values of (*p*_2_, *p*_3_) with 0 < *p*_2_ < 1 and 0 < *p*_3_ ⩽ 1, and *p*_3_ = 1 minimizes *ψ* for all values of (*p*_1_, *p*_2_) with 0 ⩽ *p*_1_ < 1 and 0 < *p*_2_ < 1. We then have *ψ*(0, *p*_2_, 1) < *ψ*(*p*_1_, *p*_2_, *p*_3_), which concludes the proof of the claim.

#### 3.2.3. 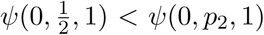

Given that we know that *p*_1_ = 0 and *p*_3_ = 1 minimize *ψ*(*p*_1_, *p*_2_, *p*_3_) at fixed *p*_2_, we have reduced this last step to a single variable problem to determine what value of *p*_2_ minimizes *ψ*(*p*_1_, *p*_2_, *p*_3_). Consider *ψ*(0, *p*_2_, 1):

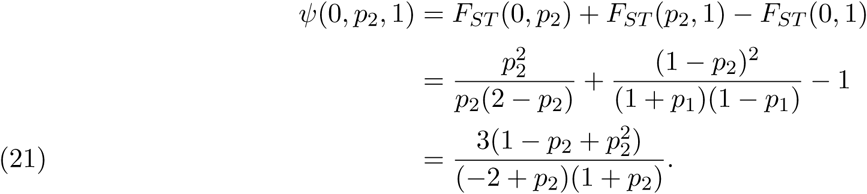

We can take the derivative of eq. (21) with respect to *p*_2_:

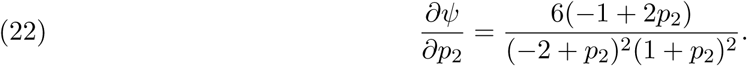

Eq. (22) is only equal to 0 when 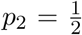. Therefore, *p*_2_ is a critical point for *ψ* in the domain 0 ⩽ *p*_2_ ⩽ 1 and specifically, 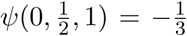 is a minimum because *ψ* is greater when *p*_2_ = 0 or *p*_2_ = 1: *ψ*(0, 0, 1) = *ψ*(0, 1, 1) = 0.

## 4. The Four-Point Condition Never Holds for Biallelic Markers with Distinct Allele Frequencies

The failure of *F*_*ST*_ to satisfy the triangle inequality for distinct allele frequencies (Section 2.3) raises the issue of the status of *F*_*ST*_ with respect to the four-point condition of Buneman (1974). The four-point condition is satisfied for a function *d* on a set *X* if and only if for all choices of four points *x*_1_, *x*_2_, *x*_3_, *x*_4_ *∈ X*, not necessarily distinct, all of the following hold:

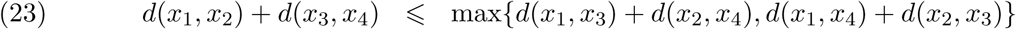

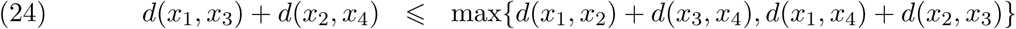

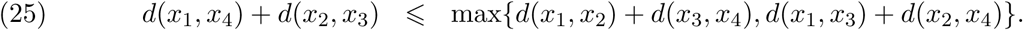

Equivalently to eqs. (23)-(25), two of the quantities *d*(*x*_1_, *x*_2_) + *d*(*x*_3_, *x*_4_), *d*(*x*_1_, *x*_3_) + *d*(*x*_2_, *x*_4_), and *d*(*x*_1_, *x*_4_) + *d*(*x*_2_, *x*_3_) are equal and greater than or equal to the third.

The four-point condition can be satisfied for some set of four points *x*_1_, *x*_2_, *x*_3_, *x*_4_ *∈ X* without necessarily holding for all sets of four points in *X*. For a specific set of four points, if and only if the four-point condition is satisfied, those points can be placed as the leaves of an unrooted tree whose edges are associated with lengths in such a manner that the pairwise distances between points computed along the tree accord with the function *d* (Buneman 1974, Steel 2016, p. 112).

For a function *d* that fails to satisfy the triangle inequality for all distinct points *x*_1_, *x*_2_, *x*_4_ in a set

*X*, the four-point condition sometimes fails; supposing *d*(*x*_1_, *x*_2_) + *d*(*x*_2_, *x*_4_) < *d*(*x*_1_, *x*_4_), we simply take *x*_3_ = *x*_2_. Noting that *d*(*x*_2_, *x*_3_) = 0, eq. (25) does not hold. However, the four-point condition can be satisfied for *x*_1_, *x*_2_, *x*_3_, *x*_4_ even if the triangle inequality is not satisfied for any three of the points, as is the case if (*d*(*x*_1_, *x*_2_), *d*(*x*_1_, *x*_3_), *d*(*x*_1_, *x*_4_), *d*(*x*_2_, *x*_3_), *d*(*x*_2_, *x*_4_), *d*(*x*_3_, *x*_4_)) = (2, 5, 8, 2, 5, 2).

We now demonstrate that for *F*_*ST*_, not only does the triangle inequality fail for all sets of three points that correspond to the allele frequencies of a biallelic marker with distinct frequencies in three populations, as shown in Section 2.3, the four-point condition also fails for all sets of four points that correspond to frequencies of a biallelic marker with distinct frequencies in four populations. This result has the consequence that sets of four populations cannot be placed on an unrooted tree in such a way that pairwise distances, as computed along the tree, accord with *F*_*ST*_.

Consider four populations whose frequencies of a particular allele at a biallelic marker satisfy 0 ⩽ *p*_1_ < *p*_2_ < *p*_3_ < *p*_4_ ⩽ 1. We show that

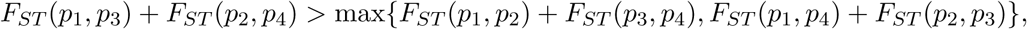

so that eq. (24) fails to be satisfied with *F*_*ST*_ in the role of *d*.

Define *s*(*p_i_, p_j_, p_k_, p_£_*) = *F_ST_* (*p_i_, p_j_*) + *F_ST_* (*p_k_, p_l_*). For *p_i_* ⩽ *p_j_* ⩽ *p_k_*, recall from eq. (12) that *ψ*(*p_i_, p_j_, p_k_*) = *F_ST_* (*p_i_, p_j_*) + *F*_*ST*_ (*p_j_, p*_*k*_) − *F*_*ST*_ (*p_i_, p*_*k*_). Applying the result of Section 2.3, *ψ*(*p_i_, p_j_, p*_*k*_) < 0 for 0 ⩽ *p_i_ < p_j_ < p*_*k*_ ⩽ 1.

We can then use eq. (12) to write

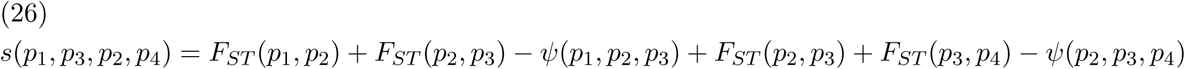

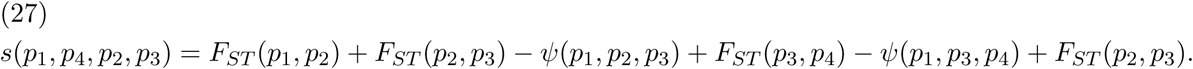

Noting that *ψ*(*p*_1_, *p*_2_, *p*_3_), *ψ*(*p*_1_, *p*_3_, *p*_4_), and *ψ*(*p*_2_, *p*_3_, *p*_4_) are all bounded above by 0 owing to the failure of the triangle inequality for *F*_*ST*_, we can cancel equal terms and use the positivity of *F*_*ST*_ in eq. (3) for distinct allele frequencies to obtain *s*(*p*_1_, *p*_2_, *p*_3_, *p*_4_) < *s*(*p*_1_, *p*_3_, *p*_2_, *p*_4_) and *s*(*p*_1_, *p*_2_, *p*_3_, *p*_4_) < *s*(*p*_1_, *p*_4_, *p*_2_, *p*_3_). We then have

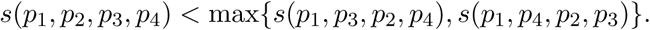

To show that the four-point condition does not hold, we must show *s*(*p*_1_, *p*_3_, *p*_2_, *p*_4_) ≠ *s*(*p*_1_, *p*_4_, *p*_2_, *p*_3_), so that with *F*_*ST*_ in the role of *d* and *p*_*i*_ in the role of *x*_*i*_, eqs. (24) and (25) cannot both hold simultaneously. Examining eqs. (26) and (27), we have

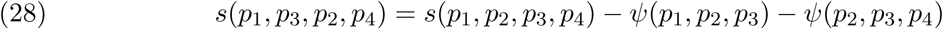

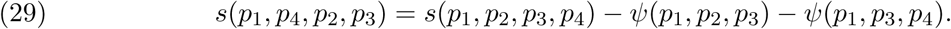

Thus, this problem reduces to showing that

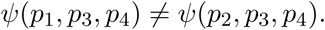

We have already shown in Section 3.2.1 that for 0 ⩽ *p_i_ < p_j_ < p_k_* < 1, *∂ψ*(*p_i_, p_j_, p*_*k*_)*/∂p_i_ >* 0.

Because *p*_1_ < *p*_2_, we can therefore conclude

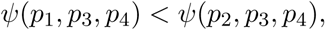

and thus, *s*(*p*_1_, *p*_3_, *p*_2_, *p*_4_) < *s*(*p*_1_, *p*_4_, *p*_2_, *p*_3_). Hence, eq. (25) fails, so that the four-point condition does not hold for distinct allele frequencies *p*_1_, *p*_2_, *p*_3_, *p*_4_. Therefore, beyond the failure of the four-point condition that results quickly when *p*_3_ = *p*_2_ from the triangle inequality not holding for 0 ⩽ *p*_1_ < *p*_2_ < *p*_4_ ⩽ 1, *F*_*ST*_ never satisfies the four-point condition when 0 ⩽ *p*_1_ < *p*_2_ < *p*_3_ < *p*_4_ ⩽ 1.

## 5. Distribution of Allele Frequencies in the Parameter Space

Next, in the context of our *F*_*ST*_ results, we consider the placement of loci from human populations in the space of possible allele frequencies. For this analysis, we examined 590,461 single-nucleotide polymorphisms (SNPs) taken from the HapMap (International HapMap 3 Consortium 2010), as used by Verdu et al. (2014) and Kang et al. (2016). We considered three populations, CEU with sample size 112 individuals, CHB with 137 individuals, and YRI with 140 individuals.

We identified ordered triples (*p*_1_, *p*_2_, *p*_3_) of frequencies, with *p*_1_ representing an allele frequency in CHB, *p*_2_ representing the frequency of the same allele in CEU, and *p*_3_ in YRI, and with *p*_1_ ⩽ *p*_2_ ⩽ *p*_3_. The SNPs can be divided into three groups based on which of the three populations has allele frequencies that lie between those of the other two populations. For the 265,517 SNPs with CEU in the intermediate position, we relabeled alleles such that 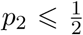. Note that we placed CEU in the intermediate position in case of ties. At nonzero allele frequencies, we observed 380 two-way ties with CHB, 621 two-way ties with YRI, and 7 three-way ties.

We plotted the values of (*p*_1_, *p*_2_, *p*_3_) over the permissible domain (Figure 2). Owing to the general similarity of allele frequencies among human populations, most points tend to have *p*_1_ only slightly less than *p*_2_ and *p*_3_ only slightly greater than *p*_2_. In regions with similar frequencies for *p*_1_, *p*_2_, and *p*_3_, *ψ*(*p*_1_, *p*_2_, *p*_3_) is only slightly less than zero. However, nontrivial numbers of points are placed in the upper left corner of the plots, where the deviation from 0 is greatest. Therefore, some SNPs in the three populations do indeed produce substantial deviations from the triangle inequality.

**Figure 2.**
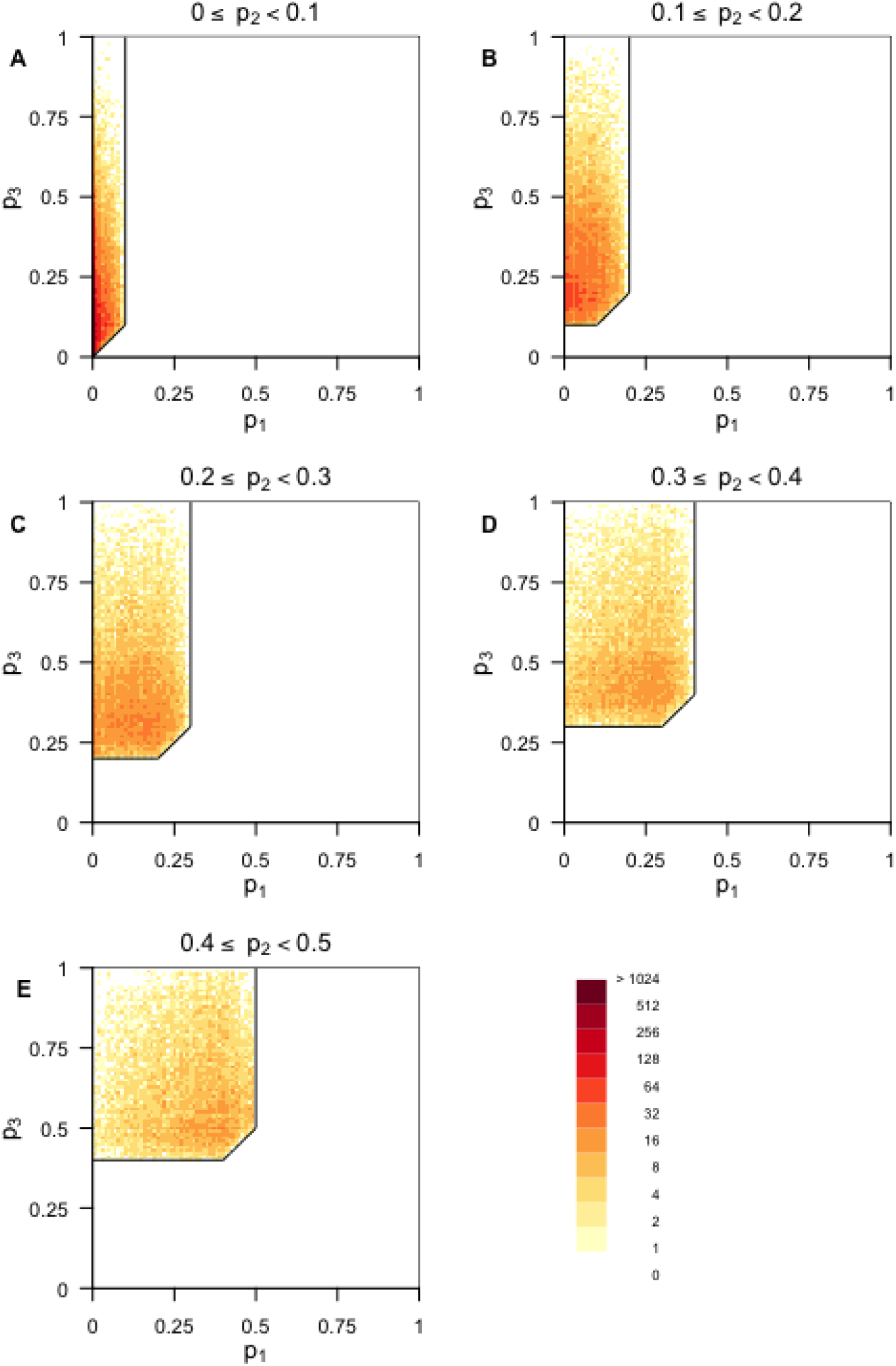
Two-dimensional histogram of population allele frequencies for three human populations: CHB (*p*_1_), CEU (*p*_2_), and YRI (*p*_3_). The plots consider 185,522 loci for which *p*_1_ ⩽ *p*_2_ ⩽ *p*_3_ when alleles are polarized by the frequency in CEU so that *p*_2_ ⩽ 0.5. (A) 0 ⩽ *p*_2_ ⩽ 0.1, (B) 0.1 < *p*_2_ ⩽ 0.2, (C) 0.2 < *p*_2_ ⩽ 0.3, (D) 0.3 < *p*_2_ ⩽ 0.4, (E) 0.4 < *p*_2_ ⩽ 0.5. The color of the box represents the number of points present, on a logarithmic scale. Because we define *p*_1_ ⩽ *p*_2_ ⩽ *p*_3_, the domain requires *p*_1_ ⩽ *p*_2_, *p*_2_ ⩽ *p*_3_, and *p*_1_ ⩽ *p*_3_. Because *p*_2_ ⩽ 0.5 is required, the upper right regions of the graph are empty. The notches in the nearly rectangular domains arise from the requirement that *p*_1_ ⩽ *p*_3_.

## 6. Discussion

In this paper, we have expanded on the observation of Sewall Wright (1978) that *F*_*ST*_ does not always satisfy the triangle inequality. In particular, we found that *F_ST_ never* satisfies the triangle inequality for biallelic markers with distinct allele frequencies. Interestingly, Wright’s case—arguably the simplest counterexample owing to its use of 0, 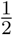, and 1 rather than more obscure frequency values—is the triplet that fails the triangle inequality by the largest amount.

### 6.1. Consequences for statistical methods

We have found that failure to satisfy the triangle inequality for all triplets of distinct allele frequencies implies failure to satisfy the four-point condition for all sets of four distinct allele frequencies. These failures to satisfy the triangle inequality and the four-point condition for all 3-tuples and 4-tuples of distinct allele frequencies for biallelic markers have implications for various forms of data analysis using *F*_*ST*_.

#### 6.1.1. Multidimensional scaling (MDS)

Matrices of pairwise dissimilarity among a set of populations are commonly used as a basis for visually depicting similarities of the populations in two or three dimensions by mulitdimensional scaling analysis (Jombart et al. 2009, Wang et al. 2010). These depictions find a representation of the matrix in a two-or three-dimensional space that has the property that Euclidean distances between points in the space approximate the matrix entries. Because *F*_*ST*_ does not satisfy the triangle inequality for biallelic markers, however, three distinct populations considered for a biallelic marker in an *F*_*ST*_ matrix cannot be represented as points in Euclidean space in such a way that Euclidean distances in the triangle connecting them correspond to the entries in the *F*_*ST*_ matrix. This imperfection of the spatial representation applies for any subset of three distinct points in a larger collection; thus, Euclidean distances between points in an MDS representation of an *F*_*ST*_ matrix necessarily only approximate the matrix entries.

Although MDS cannot always perfectly recapitulate the dissimilarities in the input matrix, MDS is frequently performed on distance matrices that are not Euclidean (Mardia et al. 1979, Cox and Cox 2001). Typical metric MDS finds a best-fit of distances between points in Euclidean space to dissimilarities in the non-Euclidean matrix. The matrix entries can also be transformed so that sets of three points necessarily satisfy the triangle inequality. One adjustment adds a constant *c* to each matrix entry (Cailliez 1983). For a dissimilarity *d*, after a large enough constant is added to obtain a new dissimilarity *d′* = *d* + *c*, the transformed distances satisfy the triangle inequality: if *d*(*x*_1_, *x*_2_) + *d*(*x*_2_, *x*_3_) < *d*(*x*_1_, *x*_3_), then a choice *c > d*(*x*_1_, *x*_3_) − *d*(*x*_1_, *x*_2_) − *d*(*x*_2_, *x*_3_) leads to *d′*(*x*_1_, *x*_2_) + *d′*(*x*_2_, *x*_3_) > *d′*(*x*_1_, *x*_3_). Alternatively, taking the square root of values in [0, 1] yields larger values still in [0, 1], so that the sum of any two of three transformed values is more likely to exceed the third one (Legendre and Legendre 1998, p. 433). In Figure 3, we apply this transformation, finding that 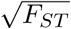 satisfies the triangle inequality for all triplets plotted.

**Figure 3.**
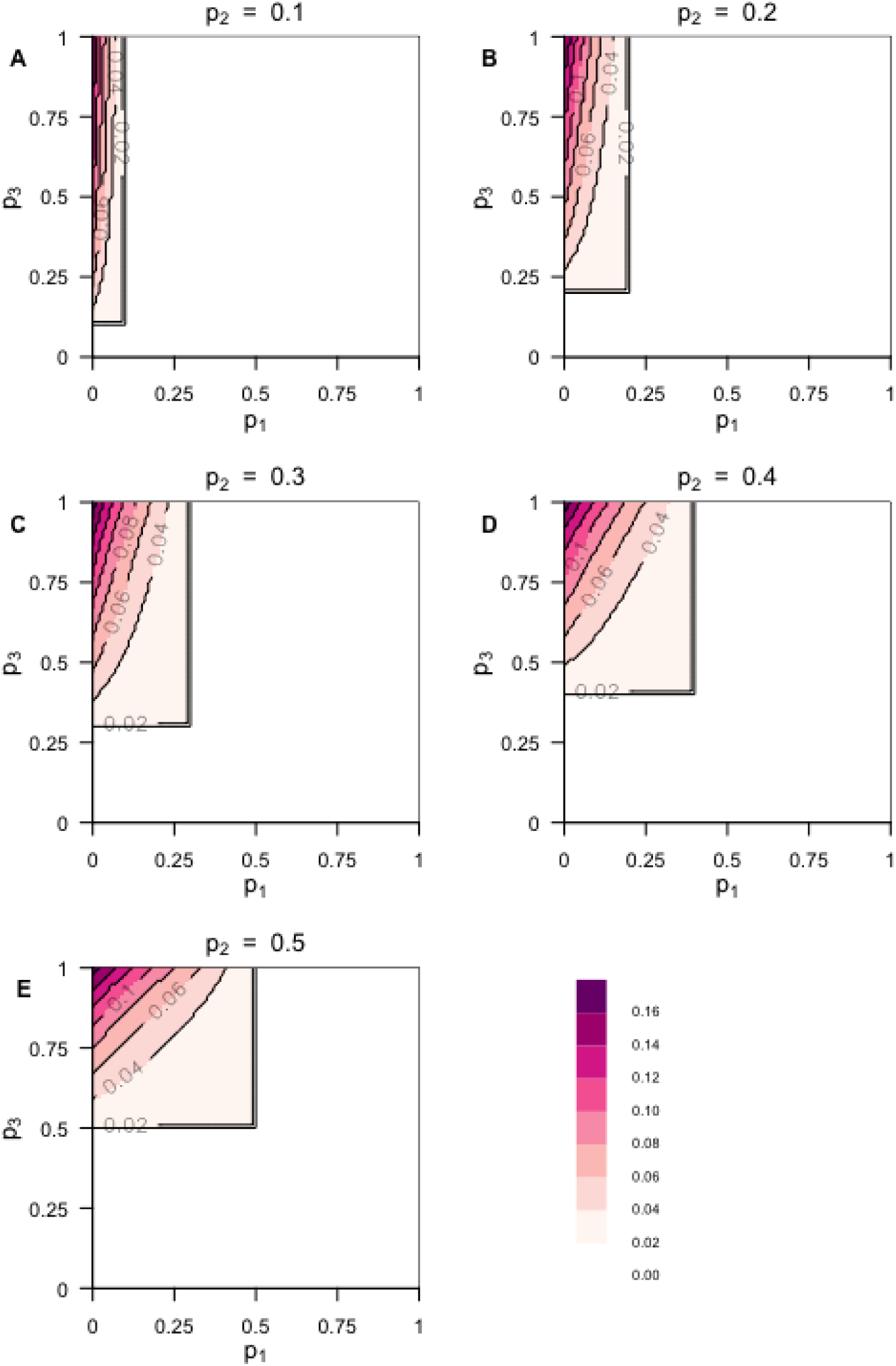
Contour plot of the relationship to the triangle inequality of the square root transformation of *F*_*ST*_. The function plotted is 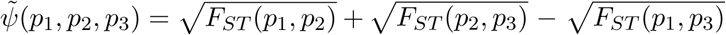 (A) *p*_2_ = 0.1. (B) *p*_2_ = 0.2. (C) *p*_2_ = 0.3. (D) *p*_2_ = 0.4. (E) *p*_2_ = 0.5. The color at a point represents the value of 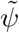 at that point. Because we define *p*_1_ ⩽ *p*_2_ ⩽ *p*_3_, the bottom and right portions of the graph are empty. Because of the symmetry of 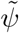 with respect to choice of allele, so that 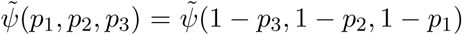 (eq. (19)), we show plots only for *p*_2_ ⩽ 0.5. For all plotted values of 0 ⩽ *p*_1_ ⩽ *p*_2_ ⩽ *p*_3_ ⩽ 1, 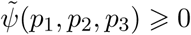.

We note, however, that the choice of transformation does affect the resulting MDS representation. In Figure 4, we compare the output of the Caillez transformation and the square root transformation on *F*_*ST*_ dissimilarity matrices with the same five allele frequencies chosen independently at random from a uniform distribution. The MDS plots differ and, in some cases, two points that are close together in one plot are distant in the other. Considering the distances between the output in the plots, neither distance matrix results in the the same matrix as the original unmodified *F*_*ST*_ dissimilarities, because *F*_*ST*_ cannot be represented as distances in Euclidean space. The choice of transform ultimately affect the results, and is relevant to report for interpretation of MDS results.

**Figure 4.**
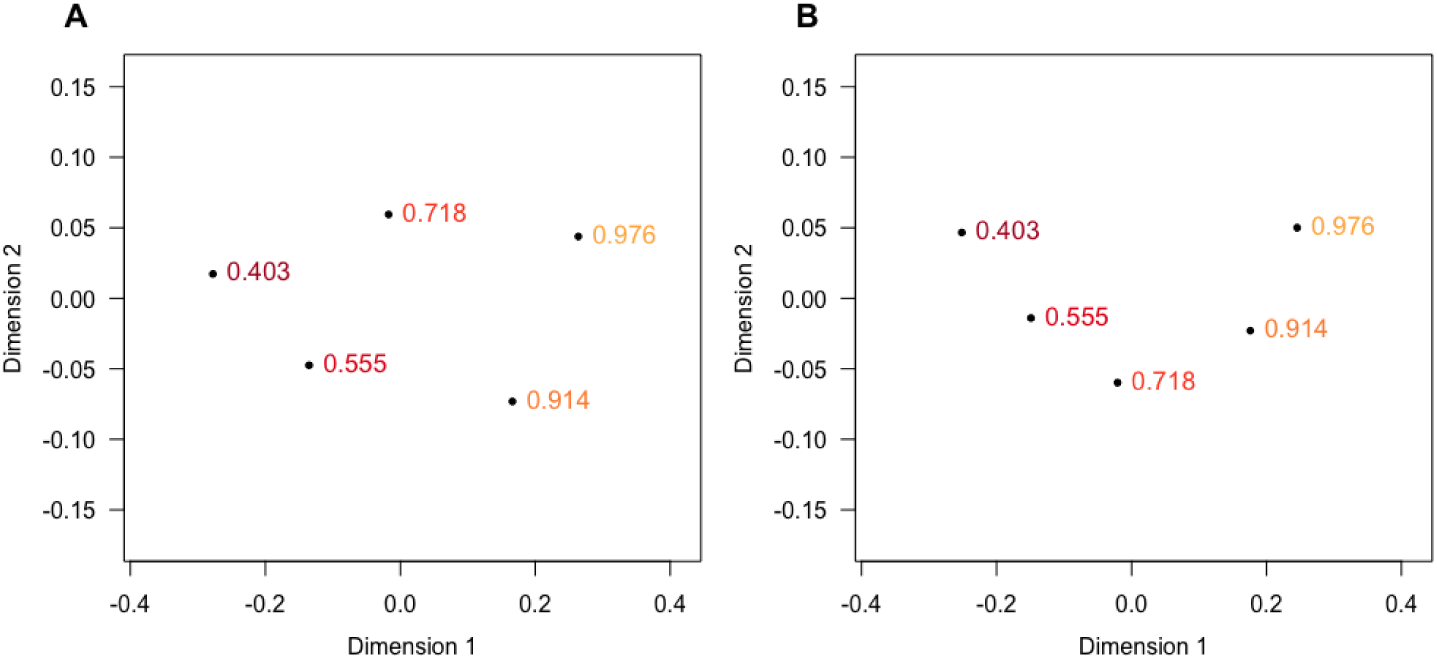
Classical multidimensional scaling (MDS) applied to transformed *F*_*ST*_ dissimilarity matrices. Five allele frequencies were chosen independently at random from a uniform-[0, 1] distribution and an *F*_*ST*_ matrix was calculated. (A) A Cailliez constant is added to all non-diagonal elements of the *F*_*ST*_ matrix. (B) The entries in the distance matrix are the square root of the pairwise *F*_*ST*_ values. The MDS output was obtained by cmdscale in R. Using the procrustes function in the R vegan package, a Procrustes analysis was used to optimally rotate the MDS output from the square-root-transformed matrix to obtain the best alignment with the output from the Caillez-transformed matrix.

#### 6.1.2. Neighbor-joining inference of evolutionary trees

A second form of analysis affected by the failure of the triangle inequality is tree reconstruction from matrices of *F*_*ST*_ values computed from allele frequencies (e.g. Takezaki and Nei 1996, Pérez-Lezaun et al. 1997, Bosch et al. 2000). Here, we consider the behavior of the neighbor-joining (NJ) algorithm applied to *F*_*ST*_ dissimilarity matrices for biallelic markers with distinct frequencies.

If a dissimilarity matrix is generated exactly from a population tree by calculating path lengths between population pairs on the tree, then NJ recovers the generating tree (Saitou and Nei 1987, Studier and Keppler 1988, Atteson 1999). Because of the failure of the four-point condition, population trees constructed from *F*_*ST*_ matrices for biallelic markers do not perfectly represent those matrices. Moreover, for any four leaves of the population tree, the minimal path connecting the leaves does not faithfully represent the *F*_*ST*_ matrix entries associated with those leaves.

Nevertheless, the inferred tree might still place more genetically similar populations close together on the tree. Using results from Mihaescu et al. (2009), we determine the tree inferred by neighbor-joining from *F*_*ST*_ matrices for 4, 5, 6, and 7 taxa. In particular, we prove the following proposition.

##### Proposition 1.

Consider a biallelic marker in n populations, with allele frequencies 0 ⩽ p_1_ < p_2_ < … < p_n_ ⩽ 1 for a specified allele. For these populations, neighbor-joining applied to an F_ST_ dissimilarity matrix, d, produces the tree topologies in Figure 5 for 4 ⩽ n ⩽ 7.

**Figure 5.**
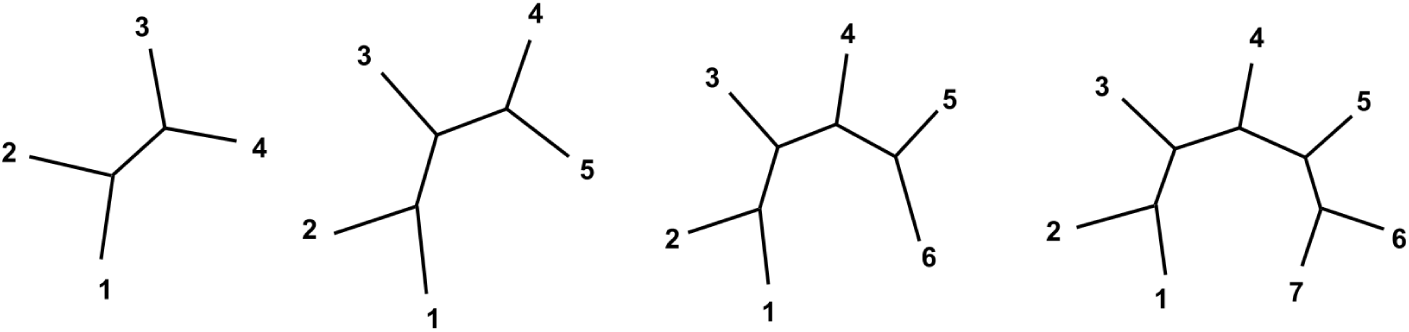
Neighbor-joining topologies for *F*_*ST*_ dissimilarity matrices with *n* = 4, 5, 6, and 7 taxa. The two cherries are (1, 2) and (*n* - 1, *n*), and all other leaves are placed sequentially along the path connecting the cherries.

The proposition states that with the populations ordered by allele frequency, neighboring populations in the sequence are placed in adjacent positions on the neighbor-joining tree. We begin with a lemma that addresses the case of *n* = 4.

##### Lemma 1.

Consider a biallelic marker in 4 populations, with allele frequencies 0 ⩽ p_1_ < p_2_ < p_3_ < p_4_ ⩽ 1. For these populations, neighbor-joining applied to an F_ST_ dissimilarity matrix, d, produces the quartet ((1,2),(3,4)).

**Proof.** Proposition 6 of Mihaescu et al. (2009) demonstrates that neighbor-joining applied to a dissimilarity *d* in four populations *i*_1_, *i*_2_, *i*_3_, *i*_4_ returns the quartet ((*i*_1_, *i*_2_), (*i*_3_, *i*_4_)) if

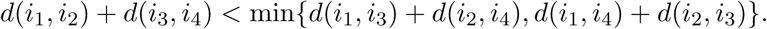

For dissimilarity measure *F*_*ST*_, we have already demonstrated in Section 4 that, using our notation, *s*(*p*_1_, *p*_2_, *p*_3_, *p*_4_) < *s*(*p*_1_, *p*_3_, *p*_2_, *p*_4_) and *s*(*p*_1_, *p*_2_, *p*_3_, *p*_4_) < *s*(*p*_1_, *p*_4_, *p*_2_, *p*_3_), so that

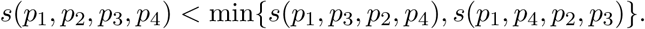

Using the definition *s*(*p_i_, p_j_, p_k_, p*_*£*_) = *F*_*ST*_ (*p_i_, p*_*j*_) + *F*_*ST*_ (*p_k_, p*_*£*_), the condition in Proposition 6 of Mihaescu et al. (2009) is obtained. □

To prove Proposition 1, we rely on the concept of *quartet consistency*, which Mihaescu et al. (2009) introduced for assessing the output topology *T* of neighbor-joining in contexts in which no tree *T* exactly captures entries in the dissimilarity matrix. By Definition 8 of Mihaescu et al. (2009), a dissimilarity map *d* for *n* populations is *quartet consistent* with a tree *T* if for every quartet ((*i, j*), (*k, l*)) *∈ T*, *w*_*d*_(*ij*: *kl*) > max[*w*_*d*_(*ik*: *jl*), *w*_*d*_(*il*: *jk*)], where

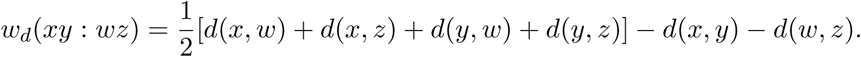

Theorem 9 of Mihaescu et al. (2009) states that for 4 ⩽ *n* ⩽ 7, if there exists a tree *T* that is *quartet consistent* with a dissimilarity map *d*: *X × X →* IR, then NJ outputs a tree with the same topology as *T*. This theorem provides a method of determining the NJ tree from a dissimilarity map on *n* taxa, 4 ⩽ *n* ⩽ 7, without proceeding through the steps of the NJ algorithm: it suffices to exhibit *T* with which *d* is quartet consistent.

##### Lemma 2.

Consider a biallelic marker in n populations, with allele frequencies 0 ⩽ p_1_ < p_2_ < … < p_n_ ⩽ 1 for a specified allele. For any i_1_, i_2_, i_3_, i_4_ with i_1_ < i_2_ < i_3_ < i_4_, using F_ST_ for the dissimilarity d, w_d_(i_1_i_2_: i_3_i_4_) > max[w_d_(i_1_i_3_: i_2_i_4_), w_d_(i_1_i_4_: i_2_i_3_)].

**Proof.** By definition of *d*, we have

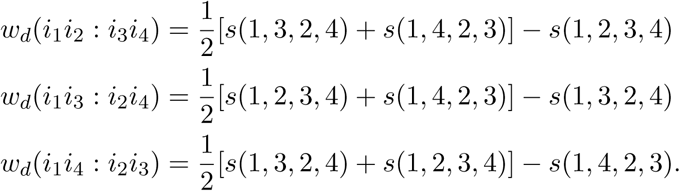

Recalling eqs. (28) and (29) and the result that *ψ*(*p*_*i*1_, *p*_*i*2_, *p*_*i*3_) < 0 for 0 ⩽ *p*_*i*1_< *p*_*i*2_< *p*_*i*3_⩽ 1,

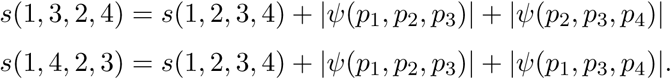

We then have:

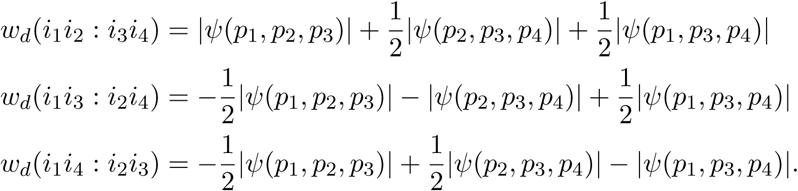

We conclude *w*_*d*_(*i*_1_*i*_2_: *i*_3_*i*_4_) > max[*w*_*d*_(*i*_1_*i*_3_: *i*_2_*i*_4_), *w*_*d*_(*i*_1_*i*_4_: *i*_2_*i*_3_)]. □

##### Proof of Proposition 1.

We consider *n* populations labeled 1, 2, *…, n* such that the frequency of a specific allele has 0 ⩽ *p*_1_ ⩽ *p*_2_ ⩽ *…* ⩽ *p*_*n*_ ⩽ 1. Consider a tree topology *T* in which populations 1 and 2 are in a cherry, *n −* 1 and *n* are in a cherry, and 3, *…, n −* 2 are arranged in numerical sequence incidental to the internal branch of the quartet ((1, 2), (*n* − 1, *n*)) (Figure 5). It suffices to show that the *F*_*ST*_ dissimilarity matrix *d* is quartet consistent with such a topology.

Consider an arbitrary subset of four populations *{i*_1_, *i*_2_, *i*_3_, *i*_4_*}*, where *p*_*i*1_< *p*_*i*2_< *p*_*i*3_< *p*_*i*4_ but *i*_1_, *i*_2_, *i*_3_, *i*_4_ are not necessarily consecutive. To show that *F*_*ST*_ is quartet consistent with *T*, it suffices to show that the quartet displayed by *T* for populations *{i*_1_, *i*_2_, *i*_3_, *i*_4_*}* has a larger value of *w* than either of the alternative quartets possible for the four populations.

By construction, *T* restricted to *{i*_1_, *i*_2_, *i*_3_, *i*_4_*}* gives quartet ((*i*_1_, *i*_2_), (*i*_3_, *i*_4_)). By Lemma 2, given *{i*_1_, *i*_2_, *i*_3_, *i*_4_*}* with *i*_1_ < *i*_2_ < *i*_3_ < *i*_4_, using *F*_*ST*_ for the dissimilarity *d*, *w*_*d*_(*i*_1_*i*_2_: *i*_3_*i*_4_) > max[*w*_*d*_(*i*_1_*i*_3_: *i*_2_*i*_4_), *w*_*d*_(*i*_1_*i*_4_: *i*_2_*i*_3_)]. Hence, *F*_*ST*_ is quartet consistent with *T*, and by Theorem 9 of Mihaescu et al. (2009), NJ applied to the *F*_*ST*_ dissimilarity matrix produces tree *T*. □

By the proposition, despite the fact that *F*_*ST*_ matrices cannot be perfectly represented on a tree, for 4 ⩽ *n* ⩽ 7 populations, NJ applied to *F*_*ST*_ places populations with neighboring allele frequencies in adjacent positions on the tree. The proof requires *n* ⩽ 7 in applying Theorem 9 of Mihaescu et al. (2009). According to that proposition, for 4 ⩽ *n* ⩽ 7, exhibiting *T* with which a dissimilarity *d* is quartet consistent suffices to determine the NJ output topology. However, for *n >* 7, quartet consistency does not suffice: it is possible to identify a tree *T* with which the dissimilarity matrix *d* is quartet consistent but that has a different topology than the tree produced by NJ.

#### 6.1.3. Estimators of F_ST_

Our results thus far have examined values of *F*_*ST*_ assuming that they are computed from true population allele frequencies. We can also examine the relationship of *F*_*ST*_ to the triangle inequality for an estimator 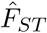.

For a biallelic locus in *K* populations with equal sample of size *n* diploid individuals, the Weir– Cockerham estimator (Weir and Cockerham 1984) is computed according to

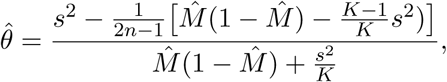

where 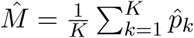 and 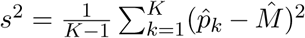 (Weir 1996, p. 173). For pairwise *F*_*ST*_, with *K* = 2, this expression simplifies to:

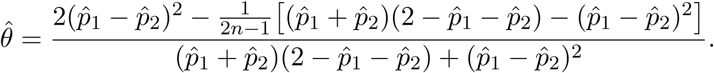

As *n → ∞*, we have

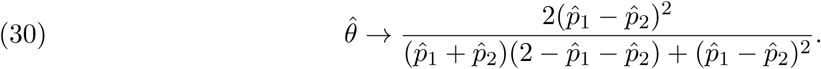

By applying the formula for *F*_*ST*_ from eq. (3), we can rewrite this limit

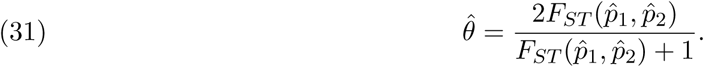

To examine whether the large-sample limit of the estimator in eq. (30) satisfies the triangle inequality, we can consider the function

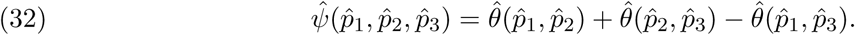

Note that 2*x/*(*x* + 1) is a monotonically increasing function for *x* in [0, 1]. Hence, using eq. (31), because *F*_*ST*_ (*p*_1_, *p*_3_) > *F*_*ST*_ (*p*_1_, *p*_2_) and *F*_*ST*_ (*p*_1_, *p*_3_) > *F*_*ST*_ (*p*_2_, *p*_3_) from eqs. (7) and (8), we have 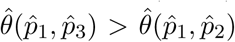 and 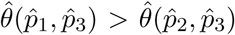. It follows that 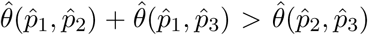 and 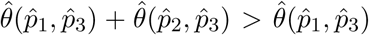. Consequently, 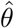 satisfies the triangle inequality for 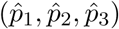 if and only if 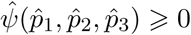

Figure 6 shows that for most of the values for 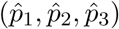 shown, 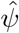 lies below zero, and hence 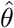 fails the triangle inequality. We did not find any values with 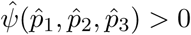. However, we did observe 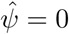 for some values with mutually distinct 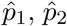, and 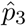, meaning that at some triples of distinct allele frequencies, 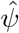 satisfies the triangle inequality. In particular, in Wright’s counterexample, 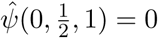. This case illustrates that the minimum of 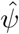 does not occur in the same place as the minimum for *ψ*. The minimum for 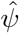 is not as far below zero as the corresponding minimum for *ψ*. From this large-sample analysis, we can conclude that the deviation from the triangle inequality is potentially not as great for the Weir–Cockerham estimator as it is for parametric *F*_*ST*_.

**Figure 6.**
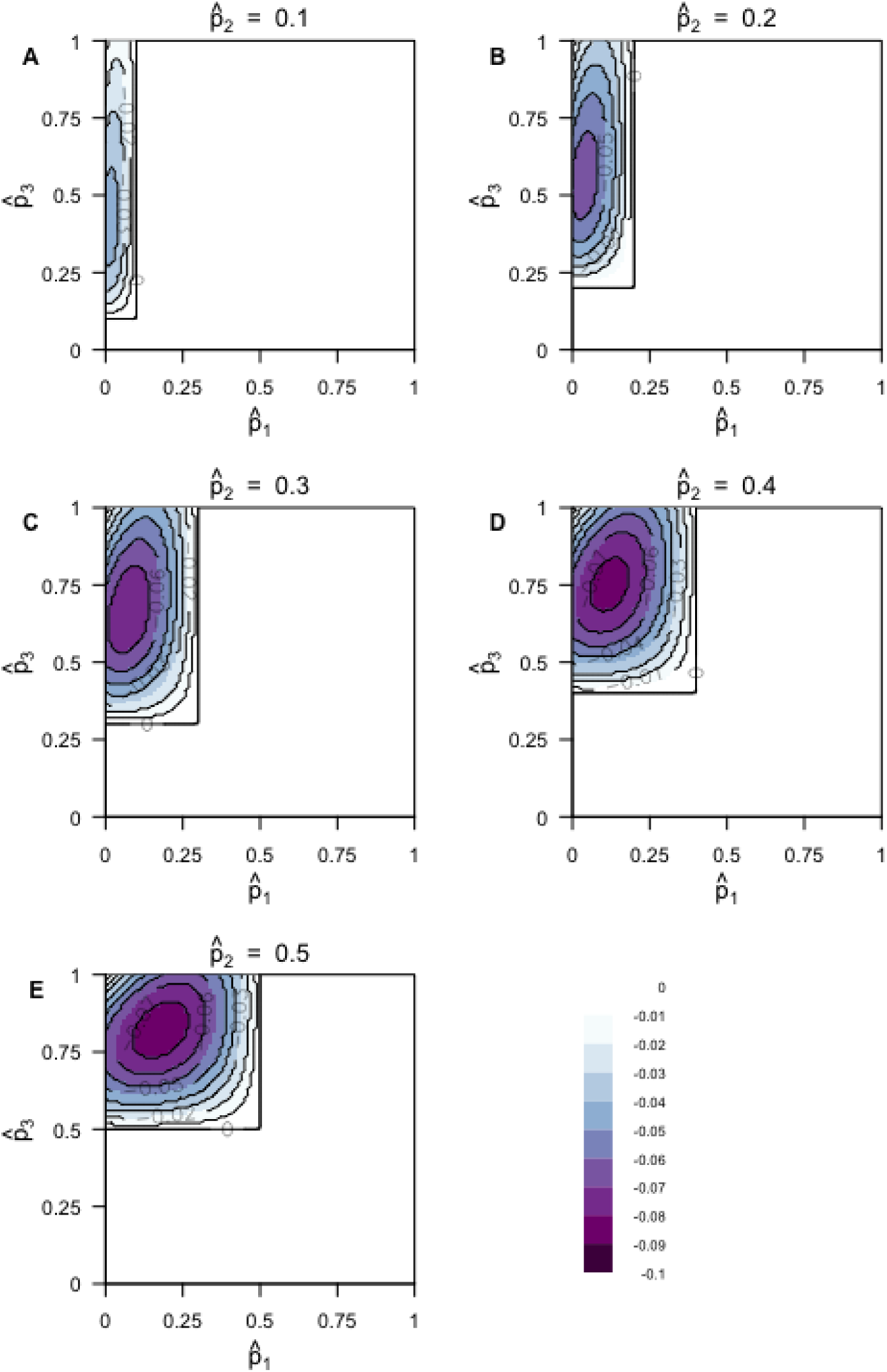
The extent to which the triangle inequality is violated for the estimator of *F*_*ST*_ as *n → ∞* (eq. (31)), 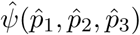, for constant 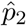 (eq. (32)). (A) 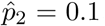. (B) 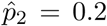. (C) 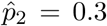.(D) 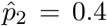.(E) 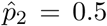. The color at a point represents the value of 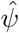 at that point. Because we define 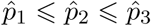>, the bottom and right portions of the graph are empty. Because of the symmetry of 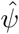 with respect to choice of allele, so that 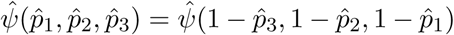(eq. (19)), we show plots only for 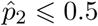.

### 6.2. Conclusions

We have seen that in allele frequencies from three human populations, the frequencies sometimes lie in parts of the allele frequency space in which the deviation is fairly large. Although we have considered sets of only small numbers of populations, relationships in a larger set of populations are constrained by features of relationships in smaller subsets, so that the results based on 3 and 4 populations that MDS and NJ representations do not perfectly represent *F*_*ST*_ matrices apply to larger sets. In the case of neighbor-joining, because demonstrating quartet consistency of a dissimilarity matrix with a tree is not sufficient to obtain the inferred tree for *n >* 7 taxa, it remains to assess the precise relationship of *F*_*ST*_ matrices and NJ.

The failure of *F*_*ST*_ to satisfy the triangle inequality can often be mitigated in data analysis. First, in the human data, the loci at which the failures are most severe correspond to points with relatively large allele frequency differences and are relatively sparse in the genome. Second, in MDS analysis, transformations can be applied to data matrices to produce spatial representations that more closely accord with the input matrix. Another solution is to use non-metric MDS, which is designed for input dissimilarity measures that are not necessarily metric; because the MDS visualization is affected by choices made in the analysis—both the version of the multidimensional scaling algorithm chosen and any associated transformations applied—it is desirable for these choices to be documented as part of the analysis (Jombart et al. 2009). Third, in NJ inference, although *F*_*ST*_ dissimilarity matrices cannot be perfectly recapitulated by a tree, we have found that NJ inference from *F*_*ST*_ produces predictable and intuitively sensible topologies for *n* ⩽ 7 taxa.

We note that we have only considered biallelic markers. Recall that the triangle inequality is satisfied for a distance between three populations if the sum of the distance between any two is greater than the third distance. Consider a modified version of *ψ* from eq. (12) for allele frequency vectors **p**, **q**, and **r** of a multiallelic marker:

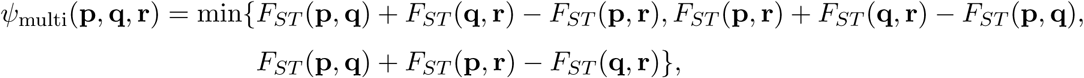

where *F*_*ST*_ follows the general eq. (1) or (2). If *ψ*_multi_ ⩽ 0, then the triangle inequality is satisfied and if *ψ*_multi_ < 0, then it fails. Taking two examples of (**p**, **q**, **r**), we have *ψ*_multi_((0, 0, 1), (1, 0, 0), (0, 1, 0)) = 1 and 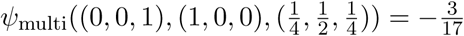. Thus, for multiallelic markers, *F*_*ST*_ sometimes satisfies the triangle inequality and sometimes does not. The multiallelic case does not have a result as simple as the biallelic result that the triangle inequality is never satisfied for distinct allele frequency vectors, and merits a more detailed analysis.

Sewall Wright’s use of a counterexample to demonstrate the failure of the triangle inequality for *F*_*ST*_ has suggested a broader investigation of the nature of *F*_*ST*_ dissimilarity matrices. The results illustrate that even fundamental statistics such as *F*_*ST*_ and simple properties such as the triangle inequality continue to permit rich mathematical analysis.

## Acknowledgments

We thank Jonathan Kang for assistance with the SNP genotypes. Support was provided by NIH grants R01 GM117590, R01 GM131404, and R01 HG005855.

